## Supplementary file for "MR-TRYX: A Mendelian randomization framework that exploits horizontal pleiotropy to infer novel causal pathways"

### Supplementary note 1: Multiple candidate traits

In practice we rely on the MR-Base database of GWAS summary datasets as a source of potential candidate traits. One outlier SNP can associate with multiple candidate traits. Sometimes this could be due to the outlier arising due to multiple independent pleiotropic pathways. Other times the same pleiotropic pathway could be represented by multiple candidate traits. For example, suppose in an analysis of exposure on outcome, a pleiotropic pathway is through adiposity. If measured lean body mass in the right arm is detected as a candidate trait, the same measurement in left arm would be redundant. It is likely for MR-TRYX to detect both as candidate traits. If, in this instance, we were to treat each of these related-candidate trait independently then we will obtain an incorrect adjustment for the outlier SNP. On the other hand, there could be instances where an outlier SNP truly does have more than one independent pleiotropic pathway to the outcome, in which case it would be appropriate to adjust for multiple candidate traits independently for that single SNP (Supplementary figure 1).

We have implemented a few practical approaches to appropriately deal with the case where multiple candidate traits associate with a single outlier.

##### Pruning

Here we estimate the adjustment independently for each candidate trait for a particular outlier and only retain the candidate trait that maximises the reduction in heterogeneity at that SNP. This approach misses the opportunity of adjusting for multiple pleiotropic pathways per outlier but is a simple approach that is less liable to misestimation or misinterpretation than the others described below.

##### Multivariable MR

If we want to make inference about Here our objective is to estimate the $(py)$ effects jointly, such that the total effect through each pleiotropic pathway is correctly estimated even if it is shared across all candidate traits. Therefore, it is not a sparse model that determines the true number of pleiotropic pathways, rather, it attempts to most accurately estimate the total effect of the pleiotropic pathways from the outlier to the outcome. The method is performed in the standard multivariable manner. First, instruments for each candidate trait are sought. Second, they are pruned into a combined instrument list that is itself clumped. Third, those SNP effects are extracted for all candidate traits and the outcome. Fourth, linear regression is performed of the SNP effects for the exposures against the SNP effects for the outcome, using the inverse of the variance of SNP-outcome estimates as weights. The regression coefficients for each exposure on the outcome are obtained from this model along with standard errors.

##### LASSO-based multivariable MR

A limitation of the standard multivariable MR approach is that it is not sparse, therefore if there are many redundant variables they will not be dropped from the model, making interpretation difficult. Here we apply shrinkage to the multivariable method using LASSO regression, with a view to drop redundant variables and therefore have a more interpretable view of the pleiotropic pathways influencing the trait. The procedure is similar to that used in the multivariable MR approach, except step 4 uses LASSO regression rather than standard linear regression. For simplicity and in the absence of an available independent dataset we use the shrinkage parameter that minimises the mean squared error. There is also an option to perform multivariable MR on only the selected parameters from the LASSO step in order to obtain effect estimates and standard errors for adjustment. This is potentially hazardous though as the standard errors are likely underestimated following the selection procedure. However, when there are a large number of candidate traits it may be necessary as multivariable MR is substantially underpowered when there are a large number of traits and the difference in SNP effects on each of them is small ^1^.

##### P-value cut-off

Here we perform multivariable MR but only retain the candidate traits that have p < 0.05.

#### Simulation 1: Accounting for redundant candidate traits

We performed simulations based on the scenario shown in Supplementary Figure 1 to evaluate the performance of the LASSO and multivariable approaches described above. The simulations involve two pleiotropic pathways from a SNP to the outcome, and each pleiotropic pathway has several redundant candidate traits. We evaluate how many candidate traits are retained in each method, the estimate of the pleiotropic pathway effect, and the precision of the estimates. In all simulations there are 50,000 samples, and each pathway explains 20% of the variance in the outcome, and 80 SNPs influencing the two pathways. In one scenario, there are 40 SNPs for pathway 1, and 40 SNPs for pathway 2. In a second scenario there are 50 SNPs influencing each pathway, with 20 of the SNPs pleiotropically influencing both pathways. In all cases each pathway has 5% of its variance explained by the SNPs that influence them. We test each of these scenarios with either both pathways having 5 redundant traits or 15 redundant traits.

The results are shown in Supplementary Table 3. Multivariable MR obtains largely unbiased estimates whereas the adjustment effects when using LASSO-based methods will be overestimated. That problem is exacerbated when there are a larger number of redundant traits. The precision, however, is very low in multivariable MR which again is exacerbated as the number of redundant variables increases. Horizontal pleiotropy does not impact any of the models substantially.

#### Simulation 2: Evaluating the sensitivity to differential instrument numbers between exposures

In this simulation, we tested if the number of instruments for the exposure influences the performance of LASSO. For example, there is a possibility that the traits instrumented by a larger number of SNPs are more likely to be retained in the model. We generated traits X_1_ and X_2_ which are instrumented by 100 genetic variants and 20 genetic variants, respectively. Each variant was bi-allelic with minor allele frequency of 0.5. Among 120 variants, 10 of them were considered as pleiotropic SNPs, having associations with X_1_ and X_2_. We considered four scenarios:

1) X1 has an effect on Y (β=0.3), where X_2_ has no effect on Y (β=0.0)
2) X1 has no effect on Y (β=0.0), where X_2_ has an effect on Y (β=0.3)
3) Both of X_1_ and X_2_ have effects on Y (β=0.3)
4) Neither of X_1_ and X_2_ have effects on Y (β=0.0)

In each scenario, we performed the LASSO regression to evaluate if the LASSO selects the traits in a manner that is determined by which trait is causal, rather than which trait has more instruments. The simulation used 10,000 individuals and was replicated 1,000 times for each scenario. The probability of how often the trait X_1_ and X_2_ are selected by LASSO were calculated and shown in Supplementary Table 4. Overall, we find that the number of instruments does not substantially influence the probability of a trait being included in the model. The LASSO kept both X_1_ and X_2_ when both traits (Scenario 3) have effects on the outcome Y with a probability of 1.000 (1000 times per 1000 simulations). Whilst, the LASSO removed both traits with a probability of 0.693 when both of X_1_ and X_2_ have no effects on Y (Scenario 4). In this scenario, the trait X_1_ was removed with a probability of 0.775 where the trait X_2_ was removed with a probability of 0.783. In the scenario 1, LASSO kept the trait X_1_ with a 100% of probability. However, LASSO failed sometime to remove X_2_, which is instrumented by a smaller number of variants, with a probability of 0.473. In scenario 2, X_2_ wasn’t removed by LASSO step but X_1_ was remained with a probability of 0.467. Considering the similar probability of the trait being eliminated in each scenario, it can be suggested that the LASSO may not favour to the candidate traits with a larger number of instruments, but it removes a trait weakly associated with the outcome.

### Supplementary figures

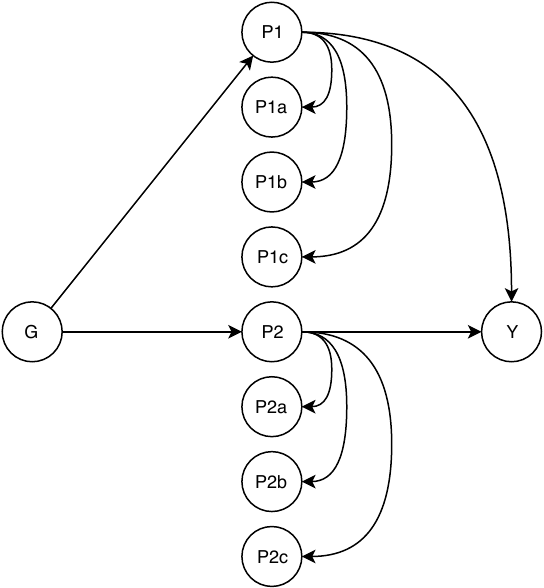

**Figure S1. The problem of multiple candidate traits associating with a single SNP in outlier adjustment.** Directed acyclic graph of the practical problem that potentially arises when searching for candidate traits associated with an outlier. Here, the outlier G has two pleiotropic pathways (P1 and P2) on the path to Y. However, P1 and P2 each have several (P1a, P1b, P1c, and P2a, P2b, P2c, respectively) alternative representations within the MR-Base GWAS summary database. Hence, it is possible that G will associate with all 8 candidate traits, when we only want representation of two pleiotropic pathways.

#### Supplementary Table S1. Sources of data used in this study.

| **Consortium/Studies** | **Use** | **Study population** | **Imputation** | **Model** | **Adjustments** |
| --- | --- | --- | --- | --- | --- |
| CARDIoGRAMplusC4D^2^ | SNP-Log odds CHD  SNP-Log odds MI | 22 233 cases and 64 762 controls of European descent  43 676 cases and 128,199 controls of European, South/East Asian, and Hispanic and African American ancestry | MACH, IMPUTE, or BIMBAM  (ref: HapMap II CEU)  IMPUTE, or MINIMAC  (ref: HapMap II/III or 1000 Genomes phase 1 ver. 3) | ADD | Age, sex and genomic control inflation factor |
| HaemGen | SNP-Platelet count^3^  SNP-Haemoglobin concentration^4^  SNP-Packed cell volume^4^  SNP-Red blood cell count^4^ | 66 867 individuals of European ancestry  71 861 individuals of European or South Asian ancestry  63 511 individuals of European or South Asian ancestry  66 214 individuals of European or South Asian ancestry | IMPUTE, BIMBAM, or MACH  (ref: HapMap II CEU)  MACH, IMPUTE, or BEAGLE  (ref: HapMap II) | ADD | Age, sex and other cohort‐specific covariates (where appropriate)  Cohort‐specific covariates  (where appropriate) |
| GUGC^5^ | SNP-Urate | 110 347 individuals of European ancestry | MACH, IMPUE, BEAGLE, BIMBAM  (ref: HapMap II CEU) | ADD | Age, sex, and study-specific covariates, if applicable |
| CKDGen^6^ | SNP-Serum Cystatin C | 133,413 individuals of European ancestry | MACH, IMPUTE, or BIMBAM  (ref: HapMap II CEU) | ADD | Age, sex, study site and genetic principal components (where appropriate). |
| IPSCSG^7^ | SNP-Primary sclerosing cholangitis | 4 796 cases and 19 955 population controls of European and American ancestry | IMPUTE2  (1000 Genomes Phase I ver. 3) | ADD |  |
| Dubois et al.^8^ | SNP-Celiac disease | 4 533 cases and 10 750 controls of European ancestry | BEAGLE and CEU, TSI, MEX and GIH  (ref: HapMap III) | ADD | Population stratification |
| GLGC^9^ | SNP-HDLC  SNP-LDLC  SNP-TC | 188 578 individuals of European, East Asian, South Asian, and African ancestry | MACH  (ref: HapMap II CEU) | ADD | Age, sex, principal components of  genomic ancestry (some studies),  and genomic control inflation factor |
| UK Biobank* | SNP-SBP  SNP-Treatments  SNP-Anthropometric measures  SNP-Smoking  SNP-Alcohol drinking  SNP-Diagnosed diseases  SNP-Education  SNP-Exercise  SNP-Memory | Approx. 340 000 individuals of European ancestry | IMPUTE2  (ref: Haplotype Reference Consortium and UK10K haplotype resource) | ADD | Population structure (where appropriate). |

CARDIoGRAM, Coronary ARtery DIsease Genome-wide Replication and Meta-analysis; GUGC, Global Urate Genetics Consortium; CKDGen, Chronic Kidney Disease Genetics; IPSCSG, International Primary Sclerosing Cholangitis Study Group; GLGC, Global Lipids Genetics Consortium; CEU, Northern Europeans from Utah; ADD, additive.

* UK Biobank GWAS results from Neale Lab: http://www.nealelab.is/blog/2017/7/19/rapid-gwas-of-thousands-of-phenotypes-for-337000-samples-in-the-uk-biobank.

**Supplementary Table S2. MR of candidate traits on each outcome (Excel).**

#### Supplementary Table S3. Results of supplementary simulation 1.

| Method | Model | Number of Redundant traits | Estimate (beta) | SE | % bias of estimate | Number of selected variables |
| --- | --- | --- | --- | --- | --- | --- |
| MVMR | Pleiotropic | 5 | 0.39 | 0.0107 | -1.89 | 12.00 |
| MVMR | Pleiotropic | 15 | 0.37 | 0.0218 | -7.02 | 32.00 |
| LASSO | Pleiotropic | 5 | 0.51 | 0.0066 | 27.30 | 6.33 |
| LASSO | Pleiotropic | 15 | 0.69 | 0.0084 | 71.49 | 10.39 |
| LASSO+MVMR | Pleiotropic | 5 | 0.53 | 0.0084 | 33.30 | 6.33 |
| LASSO+MVMR | Pleiotropic | 15 | 0.78 | 0.0120 | 95.80 | 10.39 |
| MVMR+PVAL | Pleiotropic | 5 | 0.16 | 0.0150 | -60.78 | 0.89 |
| MVMR+PVAL | Pleiotropic | 15 | 0.07 | 0.0439 | -83.52 | 2.45 |
| MVMR | Simple | 5 | 0.40 | 0.0102 | -0.79 | 12.00 |
| MVMR | Simple | 15 | 0.41 | 0.0208 | 3.18 | 32.00 |
| LASSO | Simple | 5 | 0.51 | 0.0072 | 26.39 | 6.33 |
| LASSO | Simple | 15 | 0.69 | 0.0082 | 71.37 | 10.29 |
| LASSO+MVMR | Simple | 5 | 0.53 | 0.0090 | 32.67 | 6.33 |
| LASSO+MVMR | Simple | 15 | 0.78 | 0.0116 | 95.06 | 10.29 |
| MVMR+PVAL | Simple | 5 | 0.17 | 0.0167 | -58.27 | 1.00 |
| MVMR+PVAL | Simple | 15 | 0.20 | 0.0452 | -51.24 | 2.38 |

MVMR, multivariable Mendelian randomization; LASSO+MVMR, LASSO-based multivariable Mendelian randomization; MVMR+PVAL, P-value cut-off, SE, standard error. Each row represents the results from 500 simulations. The total simulated pleiotropic effect is 0.4 across two pleiotropic pathways.

#### Supplementary Table S4. Results of supplementary simulation 2.

|  | **Probability to be selected by LASSO^1^** | | | |
| --- | --- | --- | --- | --- |
| **Scenario^2^** | **S1** | **S2** | **S3** | **S4** |
| X1 | 1.000 | 0.467 | 1.000 | 0.225 |
| X2 | 0.473 | 1.000 | 1.000 | 0.217 |

^1^ The values represent the probability of the number of times each trait is selected by LASSO out of 1000 repeated simulations. ^2^ X1 was instrumented by 100 genetic variants whilst X2 was instrumented by 20 genetic variants. ^1^ S1: X1 has an effect on Y (β=0.3), where X2 has no effect on Y (β=0.0); S2: X1 has no effect on Y (β=0.0), where X2 has an effect on Y (β=0.3); S3: Both of X1 and X2 have effects on Y (β=0.3); S4: Neither of X1 and X2 have effects on Y (β=0.0).

#### Supplementary Table S5. Proportion of unbiased estimates and AUROC value across different scenarios and methods (Excel).

**References**

1.	Sanderson E, Davey Smith G, Windmeijer F, Bowden J. An examination of multivariable Mendelian randomization in the single sample and two-sample summary data settings. *bioRxiv*, (2018).

2.	Nikpay M*, et al.* A comprehensive 1,000 Genomes-based genome-wide association meta-analysis of coronary artery disease. *Nat Genet* **47**, 1121-1130 (2015).

3.	Gieger C*, et al.* New gene functions in megakaryopoiesis and platelet formation. *Nature* **480**, 201-208 (2011).

4.	van der Harst P*, et al.* Seventy-five genetic loci influencing the human red blood cell. *Nature* **492**, 369-375 (2012).

5.	Huffman JE*, et al.* Modulation of genetic associations with serum urate levels by body-mass-index in humans. *Plos One* **10**, e0119752 (2015).

6.	Kottgen A*, et al.* New loci associated with kidney function and chronic kidney disease. *Nat Genet* **42**, 376-384 (2010).

7.	Ji SG*, et al.* Genome-wide association study of primary sclerosing cholangitis identifies new risk loci and quantifies the genetic relationship with inflammatory bowel disease. *Nat Genet* **49**, 269-273 (2017).

8.	Dubois PC*, et al.* Multiple common variants for celiac disease influencing immune gene expression. *Nat Genet* **42**, 295-302 (2010).

9.	Willer CJ*, et al.* Discovery and refinement of loci associated with lipid levels. *Nat Genet* **45**, 1274-1283 (2013).
